# Single Cell Transcriptomes, Lineage, and Differentiation of Functional Airway Microfold Cells

**DOI:** 10.1101/2023.08.06.552176

**Authors:** Manalee V. Surve, Brian Lin, Jennifer L. Reedy, Arianne J. Crossen, Anthony Xu, Bruce S. Klein, Jatin M. Vyas, Jayaraj Rajagopal

## Abstract

The airway epithelium is frequently exposed to pathogens and allergens, but the cells that are responsible for sampling these inhaled environmental agents have not been fully defined. Thus, there is a critical void in our understanding of how luminal antigens are delivered to the immune cells that drive the appropriate immune defenses against environmental assaults. In this study, we report the first single cell transcriptomes of airway Microfold (M) cells, whose gut counterparts have long been known for their antigen sampling abilities. Given their very recent discovery in the lower respiratory airways, the mechanisms governing the differentiation and functions of airway M cells are largely unknown. Here, we shed light on the pathways of airway M cell differentiation, establish their lineage, and identify a functional M cell-specific endocytic receptor, the complement receptor 2 (CR2). Lastly, we demonstrate that airway M cells can endocytose *Aspergillus fumigatus* conidia in a CR2-dependent manner. Collectively, this work lays a foundation for deepening our understanding of lung mucosal immunology and the mechanisms that drive lung immunity and tolerance.

## SHORT COMMUNICATION

The cells that sample the luminal contents of the airway remain incompletely defined. In the gut, Microfolds (M) cells have long been known to specialize in the uptake of luminal contents where they subsequently transfer their endocytosed cargo to the immune cells of gut-associated lymphoid tissue (GALT), classically known as Peyer’s patches^1-4^. In the upper airway, M cells are located adjacent to nasal-associated lymphoid tissues (NALT),^5-7^ but their presence in the normal lung remained uncertain. Then, in 2019, murine inflammatory models were used to definitively demonstrate that lung M cells existed^8^. Given the very recent discovery of airway epithelial M cells, the key signalling pathways governing their differentiation and developmental origins have not been delineated.

Although murine airway inflammation clearly resulted in the differentiation of abundant airway M cells, we reasoned that homeostatic M cells might have escaped attention due to their scarcity, as was the case with the elusive pulmonary ionocyte^9^. Therefore, we reanalyzed our prior single cell transcriptomic data containing 69,607 murine tracheal epithelial cells^9^ and probed for canonical M cell markers. We uncovered a cluster of epithelial cells that we had previously misannotated. This cluster is characterized by the expression of classical M cell markers including the *Spib* and *Sox8* transcription factors, the chemokines *Ccl9* and *Ccl20*, and *Tnfrsf11a* (the Receptor activator of NF-*κ*B; RANK) which is known to be involved in gut M cell differentiation^10^ (Figure 1A and B, Figure E1A, and Table E1). We also identified the bacterial uptake receptor *Gp2*^*4*^, and a host of genes associated with gut M cells including *Tnfaip2, Marcksl1, and Anxa5* (Figure 1B, Figure E1A, and Table E1)^1^. We previously miscalled these M cells as immune cells because they had been clustered together with immune cell populations due to their shared chemokine expression. Indeed, we only identified 58 M cells out of 69,607 airway epithelial cells, making them the single most scarce population in the airway epithelium (0.08%) even when compared to ionocytes, neuroendocrine cells, and tuft cells (Figure E1B)^9^.

**Figure 1.**
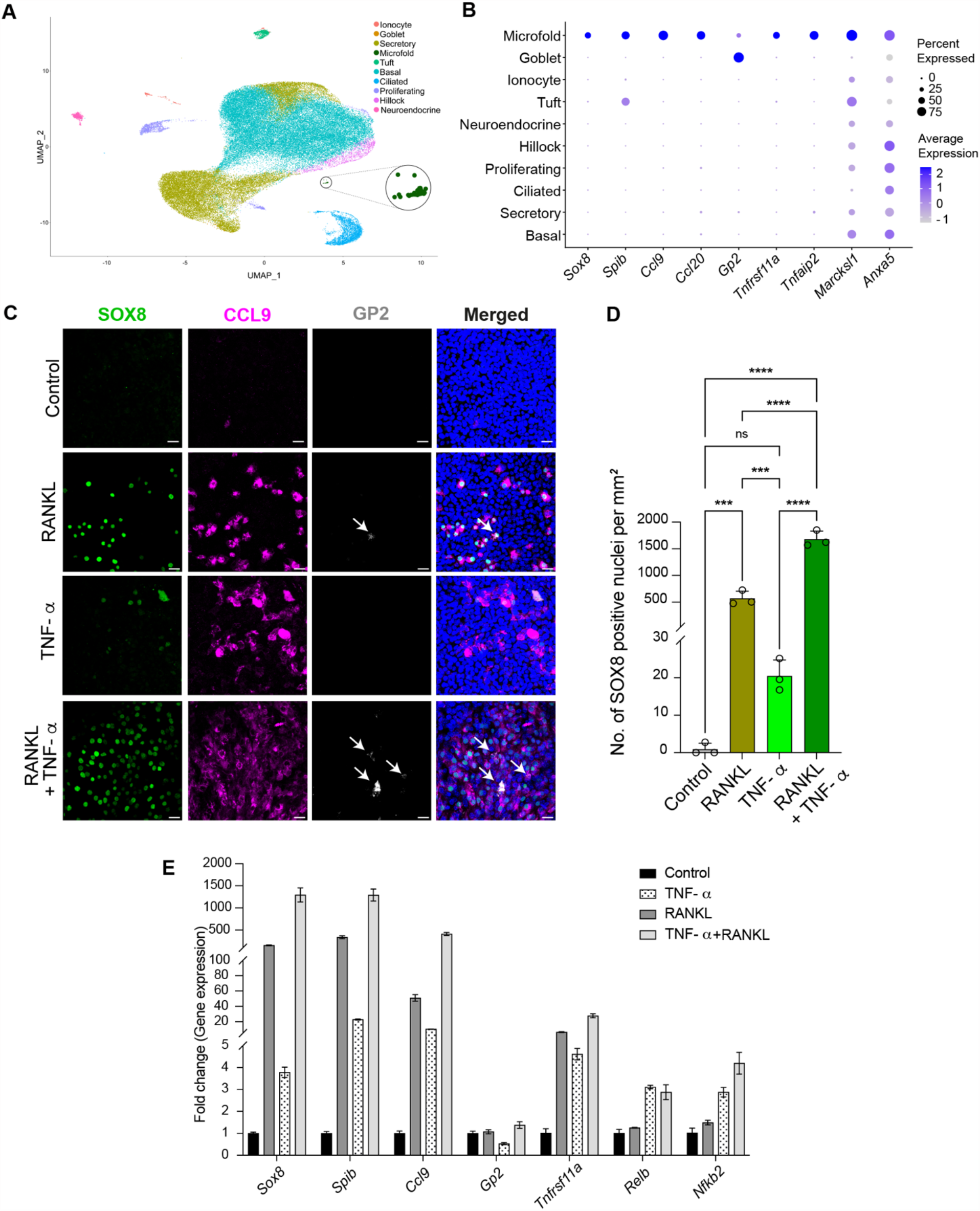
M cell transcriptomes and mechanisms of *in vitro* differentiation. *(A)* Dimensionality reduced visualization using Uniform Manifold Approximation and Projection (UMAP) of scRNA-seq transcriptomes of tracheal epithelium showing a newly identified M cell cluster (circled and enlarged). *(B)* Dot plot showing that M cells express characteristic M cell transcription factors, chemokines, receptors, and other M cell markers. *(C-E)* Mature ALI epithelial cultures derived from primary murine basal stem cells from the trachea were treated daily for 6 days with RANKL (200 ng/ml), TNF-*α* (50 ng/ml), or combined RANKL and TNF-*α* for 6 days to induce M cells. PBS served as controls. *(C)* Confocal micrographs of representative regions from mouse ALI epithelial cultures immunostained with characteristic M cell markers, SOX8 (green), CCL9 (magenta), and GP2 (gray). Nuclei are stained with Hoechst dye (blue). GP2 positive mature M cells are marked with white arrows. Scale bar, 20 *μ*m. *(D)* Quantification of SOX8 positive nuclei following indicated treatments. Individual circles on the graph represent the percent of all epithelial cells that are SOX8 positive M cells in single ALI epithelial culture wells treated with the indicated cytokines. Data are presented as mean ± SD (standard deviation) of triplicate ALI cultures. ns = not significant, \*\*\**P* ≤ 0.001, and ****P≤ 0.0001 by one-way ANOVA. *(E)* qPCR-based expression analysis of ALI epithelial cultures for the indicated M cell specific and non-canonical NF-*κ*B pathway genes following the specified treatment conditions. Data are presented as mean ± SD of triplicate ALI epithelial cultures.

In the gut, RANK signaling through the non-canonical NF-*κ*B pathway is sufficient to induce M cell differentiation from Lgr5 positive intestinal stem cells^10, 11^. Indeed, airway M cells were first identified following RANKL (RANK ligand) induction^6, 8^. In the gut, TNFα-induced canonical NF-*κ*B pathway signaling cooperates with RANKL-mediated non-canonical NF-*κ*B pathway activation to further increase M cell differentiation^12^. Reasoning that these pathways of differentiation would be conserved in gut and airway, we hypothesized that TNF-α might enhance RANKL-induced airway M cell differentiation. Therefore, we treated murine tracheal air-liquid interface (ALI) epithelial cultures with RANKL or TNF-α alone or in combination, and then performed immunohistochemistry for the canonical M cell markers, SOX8, CCL9, and the mature M cell marker GP2^13^. As expected, RANKL, induced M cells (Figure 1C and D)^8^. Surprisingly, unlike the case in the gut, the administration of TNF-*α* alone was capable of inducing a small number (∼20 SOX8 positive M cells/mm^2^) of GP2 negative immature M cells (Figure 1C and D)^13^. However, M cell differentiation was the greatest in cultures exposed to combined TNF-*α*/RANKL treatment with a 2.9-fold higher M cell induction when compared to RANKL stimulation alone (Figure 1C and D). Additionally, combined TNF-*α*/RANKL treatment induced higher expression of the M cell genes *Sox8, Spib, Ccl9*, and *Gp2* when compared to RANKL treatment alone (Figure 1E). Notably, TNF-*α*, which acts through the canonical NF-*κ*B pathway, upregulates the expression of *Tnfrsf11a* (the RANK receptor), and the non-canonical NF-*κ*B monomer genes, *RelB* and *Nfkb2* (p52) in addition to M cell-specific genes (Figure 1E). Collectively, these results suggest that TNF-*α*, in addition to augmenting RANKL-mediated M cell differentiation as it does in the gut, can by itself upregulate M cell genes and induce the differentiation of immature airway M cells.

In our scRNA-seq datasets, we observed that a fraction of both basal stem cells and secretory cells expresses *Tnfrsf11a* transcripts, suggesting the hypothesis that both these cell types might be able to differentiate into M cells (Figure 2A). Therefore, we established ALI epithelial cultures from *p63CreER::tdTomato*^*fl/fl*^ and *Scgb1a1CreER::tdTomato*^*fl/fl*^ transgenic mice which were used to demonstrate whether M cell progeny arise from parent basal and/or secretory cells, respectively. Following lineage labeling, ALI cultures were treated with RANKL to induce M cell differentiation (Figure 2B). Although SOX8 positive M cells did originate from basal stem cells, M cells were considerably more likely to arise from secretory cells (Figure 2C and D). We note that goblet cells and ciliated cells also predominantly originate from secretory cell parents during homeostasis, but both goblet and ciliated cells can directly differentiate from basal stem cells in the setting of injury^14^.

**Figure 2.**
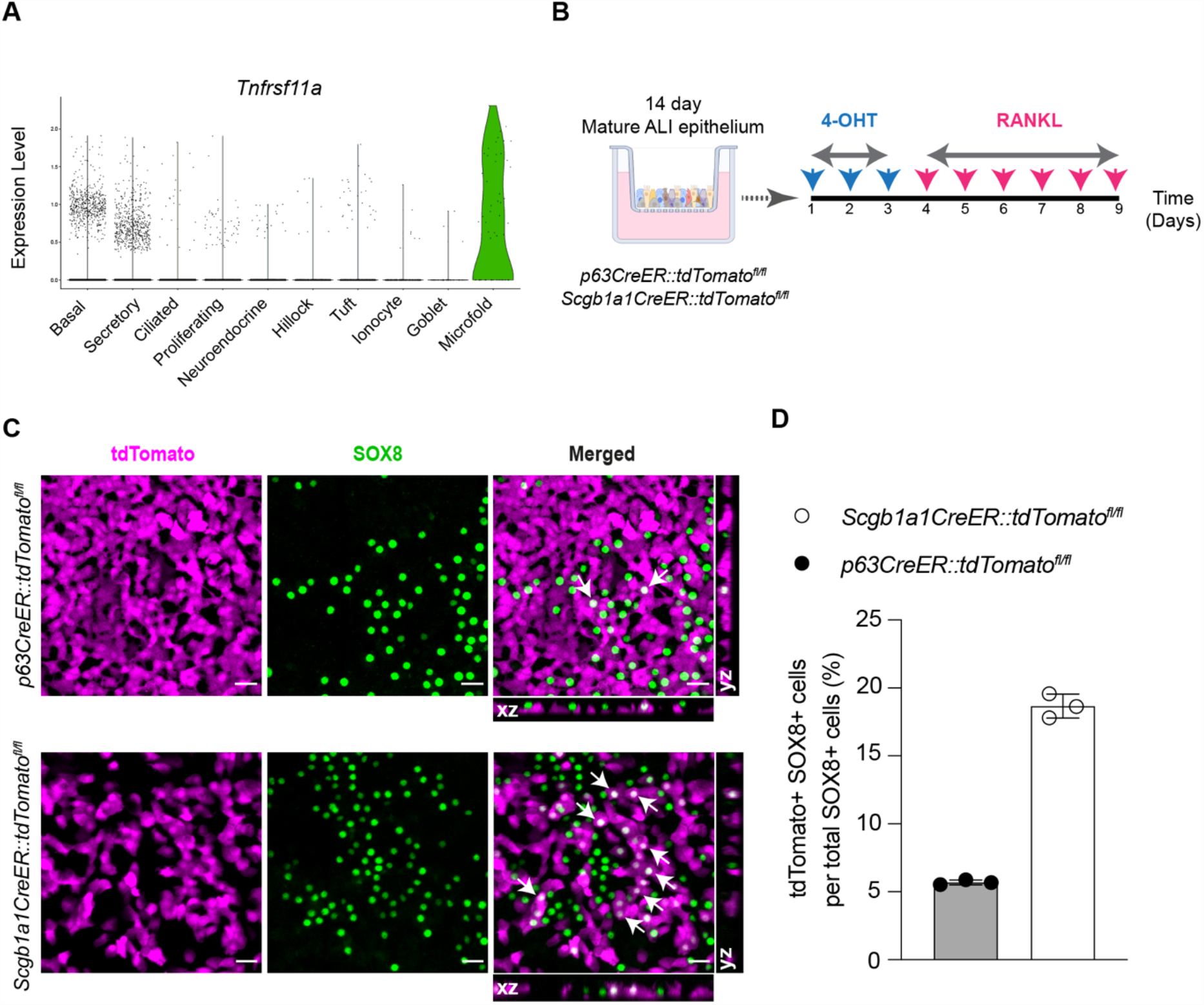
*In vitro* lineage tracing of airway M cells. *(A)* Violin plot of scRNA-seq profiles showing expression of *Tnfrsf11a* gene (RANK) in M cells as well as a small fraction of airway basal stem and secretory cells. *(B)* Schematic of *in vitro* lineage tracing of M cells in mouse tracheal ALI epithelial cultures derived from mice expressing cell-specific CreER drivers for basal stem (*p63CreER*) and secretory (*Scgb1a1CreER*) cells and the LSL-tdTomato reporter. 4-OHT stands for 4-hydroxytamoxifen. *(C)* Immunofluorescence images of *p63CreER::tdTomato*^*fl/fl*^ and *Scgb1a1CreER::tdTomato*^*fl/fl*^ lineage traced epithelial cells following RANKL-mediated M cell induction. Lineage positive M cells marked with arrows showing cytoplasmic tdTomato (magenta) and nuclear SOX8 (green) staining are seen in the merged panels and orthogonal sectional views. Scale bar, 20 *μ*m. *(D)* Quantitative analysis of percent lineage positive M cells. Individual circles on the graph represent lineage positive cells in single RANKL-treated ALI epithelial culture wells. Data are presented as mean ± SD of triplicate ALI epithelial cultures.

We next sought to address whether airway M cells possess the machinery for the uptake of luminal cargo. Interestingly, *Cr2*, encoding for a complement receptor, is expressed in a subset of airway M cells (Figure 3A and E1A) and we confirmed protein expression using confocal microscopy (Figure 3B). It is well known that complement deposition on *Aspergillus* conidia facilitates pathogen clearance^15^. Therefore, we hypothesized that airway M cell complement receptor 2 (CR2) would enhance the uptake of *A. fumigatus* conidia. To determine the role of CR2 in M cell-mediated uptake of *Aspergillus fumigatus (Af)*, we performed antibody-mediated receptor blockade in RANKL-treated ALI epithelial cultures. Subsequently, we exposed these cultures to *Aspergillus* conidia that were pretreated with PBS or complement containing serum. To quantify intracellular conidia, we used a fluorescent *Aspergillus* reporter (FLARE) strain that expresses DsRed. We observed *Af* conidia being endocytosed specifically by M cells (CCL9^+^ cells) only in RANKL-treated cultures (Figure 3C). Additionally, we observed a dramatic reduction in the uptake of complement-coated *Af* conidia following CR2 antibody-neutralization as compared to the IgG1 control (Figure 3C and D). This effect was confirmed using epithelia generated from CR2^-/-^ mice (Figure 3E).

**Figure 3.**
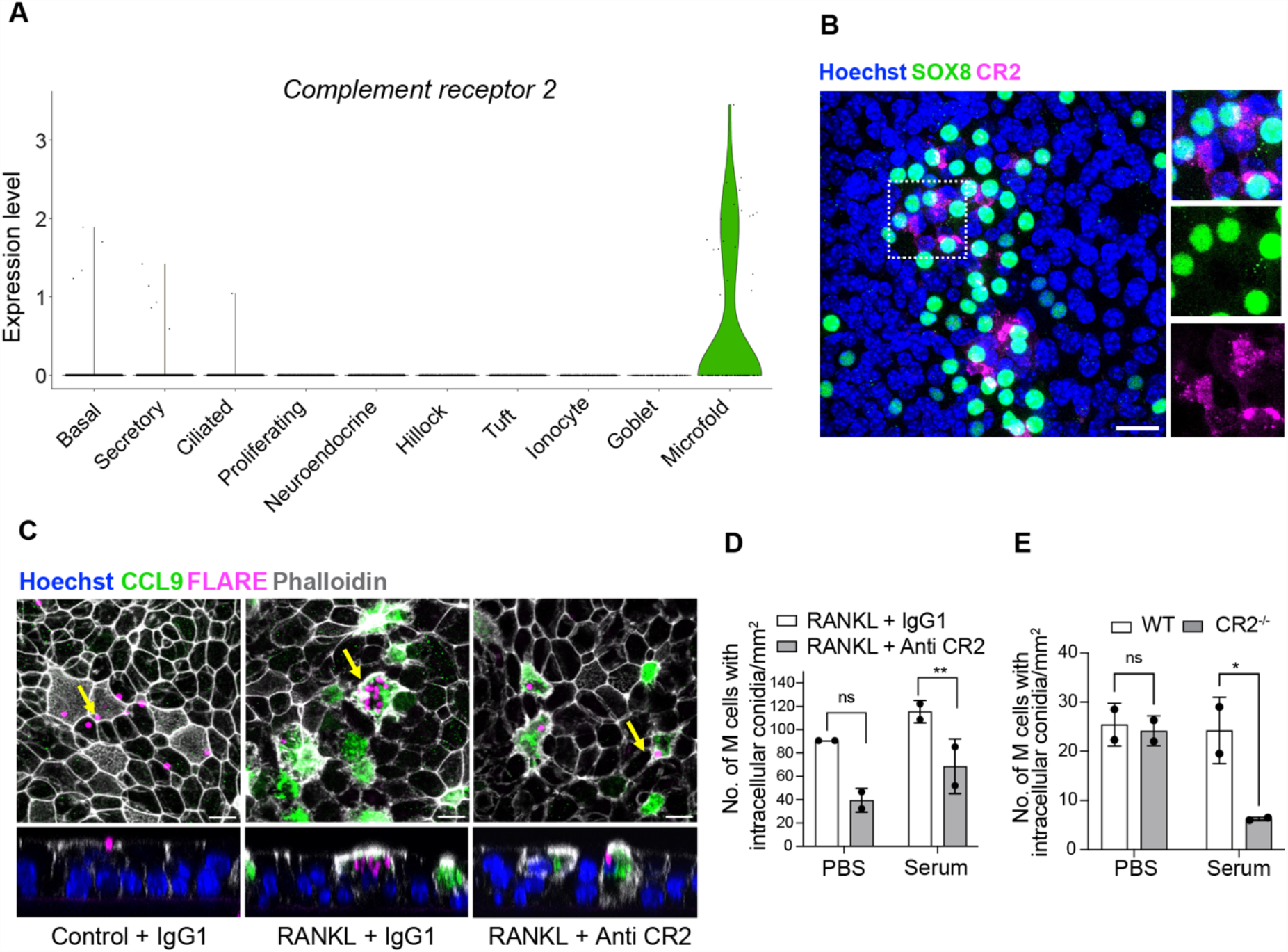
Airway M cells express complement receptor 2 (CR2) and internalize *Aspergillus fumigatus* conidia in a CR2-dependent manner. *(A)* Violin plot showing specific expression of *Cr2* in M cells based on scRNA-seq profiles. *(B)* Confocal micrograph of RANKL-induced mouse ALI epithelial cultures stained for M cell marker SOX8 (green) and CR2 (magenta) demonstrates CR2 expression in a subset of M cells. Boxed region is enlarged in insets on the right. Nuclei are stained with Hoechst dye (blue). Scale bar, 20 *μ*m. *(C)* Confocal images showing FLARE conidia (magenta) adhering extracellularly to epithelial cells in control ALI (PBS) (left), internalization of multiple conidia inside CCL9 positive M cells (green) in RANKL-treated ALI (middle) and uptake of fewer conidia by M cells in RANKL-treated ALI pre-treated with Anti-CR2 antibody (Ab) (right). Epithelial cell membranes are marked by phalloidin (white). The panels below are orthogonal views (YZ) that include the cells marked by arrows in the respective panels above. Nuclei are stained with Hoechst dye (blue). Scale bar, 10 *μ*m. *(D)* Quantification of endocytosed intra-M cell complement coated or uncoated FLARE conidia in RANKL-treated ALI cultures pre-treated with Control IgG1 or anti-CR2 antibodies. *(E)* Quantification of complement coated or uncoated FLARE conidia within M cells in RANKL-treated ALI cultures derived from wildtype (WT) or CR2^-/-^ (CR2 knockout) basal cell lines. Individual circles on the graph represent intra-M cell conidia in single RANKL-treated ALI epithelial culture wells. Data are presented as mean ± SD of duplicate ALI cultures. ns = not significant, \*\**P* ≤ 0.01, and \**P* ≤ 0.05 by two-way ANOVA (Tukey’s multiple comparisons test).

In summary, we report the first single cell transcriptomes of airway M cells, demonstrate conservation in the mechanisms of M cell differentiation in the gut and airway, identify the lineage of airway M cells, and report a model for functionally interrogating airway M cell uptake.

## ACKNOWLEDGMENTS

We would like to thank Drs. Avinash Waghray, Jiajie Xu, Jiawei Sun, Qiaozhen Liu, Gergana Shipkovenska, Viral Shah, Chaim Chernoff, and other members of the Rajagopal and Klein Labs for their valuable feedback. This research was supported by NIH project number 1R01HL164563 and 1K08AI141755.

