## Supplementary Information for "Single Cell Transcriptomes, Lineage, and Differentiation of Functional Airway Microfold Cells"

### MATERIALS AND METHODS

#### Single cell RNA sequencing analysis

Existing scRNA-seq data was accessed from GEO accession GSE103354, previously mapped to MM10 using Cellranger v1.0.1. The RDS file was imported into R v4.1.3 and processed as a Seurat object using Seurat v4.3.0. Cells with less than 1000 detected genes or greater than 5% mitochondrial read counts were filtered. FindClusters with a resolution of 0.5 detected a split cluster marked by immune genes such as *Ptprc* (CD45). Sub-clustering on this population resolved a distinct cluster that was *Ptprc* negative and *Epcam* positive, which was later identified as a cluster of M cells.

#### Mouse models

C57BL/6J (stock no. 000664), $CR2^{-/-}$ (stock no. 008225), and tdTomato mice (stock no. 007914), were purchased from Jackson laboratory. *p63-CreER*[1] and *Scgb1a1-CreER*[2], mice were previously described. Mice were maintained in an Association for Assessment and Accreditation of Laboratory Animal Care-accredited (AAALAC) animal facility at the Massachusetts General Hospital. All procedures were performed with Institutional Animal Care and Use Committee (IACUC)-approved protocols. All mice were housed in an environment with controlled temperature and humidity, on 12-hour light:dark cycles, and fed with regular rodent's chow.

#### Air-liquid Interface epithelial model for M cell induction

Murine tracheal cells were harvested and cultured as previously described[3]. Following enzymatic dissociation (papain) of trachea, basal stem cells were cultured and expanded in complete SAGM (small airway epithelial cell growth medium; Lonza, CC-3118) containing a TGF-β/BMP4/WNT antagonist cocktail and 5 μM ROCK inhibitor Y-27632 (Selleckbio, S1049). Airway basal stem cells were trypsinzed and seeded onto 0.4 μm transwell membranes pre-coated with 804G-conditioned medium and grown in SAGM media for 24 h. The following day, air-liquid interface (ALI) epithelial cultures were initiated by aspirating the media and adding PneumaCult-ALI Medium (Stem Cell Technologies, 05001) to the bottom chamber only. Differentiation of airway basal stem cells on an ALI was followed by directly visualizing beating cilia in real time after 14 days. To induce M cells, 14-day mature ALI epithelial cultures were treated daily with 200 ng/ml of RANKL or 50 ng/ml of TNF-$\alpha$ alone (Peprotech) or in combination in PneumaCult-ALI Medium for 6 days.

#### *In vitro* lineage tracing

To perform *in vitro* lineage tracing, basal stem cells derived from trachea of *p63CreER::tdTomato*$^{fl/fl}$ and *Scgb1a1CreER::tdTomato*$^{fl/fl}$ transgenic mice were used to establish ALI epithelial cultures as above. To label parent basal stem or secretory club cells, 14-day mature ALI epithelial cultures were treated for 3 days with 1 $\mu$m of 4-hydroxytamoxifen (Sigma-Millipore). Subsequently, ALI cultures were treated with 200 ng/ml of RANKL daily for 6 days to induce M cells.

#### Fungal culture

The Fluorescent Aspergillus Reporter (FLARE) strain[4], a derivative of the CEA10 strain of *Aspergillus fumigatus* was grown at 37°C for 3 days on glucose minimal media slants. Slants were seeded from frozen stocks of conidia maintained at −80°C. To harvest conidia, a sterile solution of deionized water (Milli-Q; Millipore Sigma, Burlington, MA) containing 0.01% Tween 20 was added to each slant, and spores were liberated using gentle surface agitation with a sterile swab. The spore solution was passed through a 40-µm cell strainer to separate hyphal debris. Spores were washed three times with sterile PBS and counted on a LUNA automated cell counter (Logos Biosystems, Annandale, VA) and diluted appropriately.

#### Complement receptor blockade/knockout and fungal uptake assay

To determine the role of complement receptor 2 (CR2) in M cell uptake of *Aspergillus fumigatus*, we performed antibody-mediated receptor blockade. Briefly, control and RANKL-treated ALI epithelial cultures were pretreated with IgG1 control antibody (10 $\mu$g/ml, BioXcell) or anti-CR2 antibody (10 $\mu$g/ml, Abcam-Clone 4B2) for 2 h. Subsequently, ALI epithelial cultures were apically exposed to PBS or serum pre-treated (2 h, to allow complement deposition) FLARE conidia at a concentration of 1 x $10^7$ conidia/$cm^2$ for 4 h. In another experiment, tracheal basal stem cells derived from Wildtype C57BL/6J and $CR2^{-/-}$ mice were differentiated into mature ALI epithelial cultures and treated with PBS or RANKL (200ng/ml, 6 days) to induce M cells. Next, ALI epithelial cultures were infected with PBS or serum pretreated FLARE conidia for 4 h. Subsequently, ALI epithelial cultures were immunostained and imaged using confocal microscopy to quantify the numbers of intracellular conidia.

**Immunostaining, microscopy, and image analysis**

The following antibodies were used: SOX8 (1:500, sc-374445, Santacruz Biotechnology), CCL9 (1:500, AF463, R & D Biosystems), GP2 (1:500, D278-3, MBL), and Alexa Fluor 594 conjugated CR2 (1:100, 123426, Biolegend). All Donkey Alexa Fluor conjugated secondary antibodies (488, 594 and 647) were obtained from Thermo Fisher Scientific and were used at 1:500 dilution. F-actin filaments were stained using Alexa Fluor 647 conjugated Phalloidin (1:200, A22287, Thermo Fisher Scientific).

ALI epithelial cultures were fixed in 4% PFA for 10 mins at room temperature and washed with PBS. Epithelial cultures were then incubated with primary antibodies in staining buffer (0.1% Triton X-100 and 1% BSA in PBS) overnight at 4 °C on an orbital shaker. The following day, cultures were washed with PBS, incubated with appropriate secondary antibody in staining buffer for 1 h at room temperature, washed with PBS, and counterstained with Hoechst 33342. Finally, ALI epithelial cultures were mounted on glass slides using Fluoromount-G mounting medium (SouthernBiotech).

Microscopic images were obtained with an Olympus FV10i confocal laser-scanning microscope or with a Leica SP8 confocal laser-scanning microscope using a 60x oil and 40x water objective, respectively. Image processing and analysis were done using Fiji software.

**Statistical analysis**

All statistical analyses were done using GraphPad Prism software (version 10.0.1). Statistical test applied to each experimental paradigm and the accepted statistical significance values are stated in the respective figure legends.

### SUPPLEMENTAL TABLE

**Table E1:** List of the top 200 differentially expressed genes in airway M cells. Within each column of the table, 'gene' represents the gene symbol, 'pct.1' represents the percentage of cells where a gene is detected in the M cell cluster ($1^{st}$ group), 'pct.2' represents the percentage of cells where a gene is detected in other clusters ($2^{nd}$ group), 'avg_Log2FC' represents log fold-change of the average expression between $1^{st}$ and $2^{nd}$ groups, p_val and p_val_adj represent p-value and adjusted p-value, respectively.

### SUPPLEMENTAL FIGURE

**Figure E1.** Single cell transcriptomic analysis of M cells in murine tracheal epithelium. *(A)* Heat map of airway M cell genes that were differentially expressed relative to the genes of other airway epithelial cell types. Note that to match the number of cells in the M cell cluster, we considered top 50 cells from other epithelial cell clusters to generate the heatmap. *(B)* Percentage of cell type abundance in the scRNA-seq profiles of murine tracheal epithelium.

**Table E1**

| gene | pct.1 | pct.2 | avg_log2FC | p_val | p_val_adj |
|---|---|---|---|---|---|
| *Ccl9* | 0.845 | 0.011 | 5.436869 | 0 | 0 |
| *Ubd* | 0.69 | 0.003 | 3.622734 | 0 | 0 |
| *Spib* | 0.69 | 0.008 | 3.54316 | 0 | 0 |
| *Csn2* | 0.483 | 0 | 3.190641 | 0 | 0 |
| *Wfdc18* | 0.483 | 0.006 | 2.989371 | 0 | 0 |
| *Bcl2a1b* | 0.759 | 0 | 2.785754 | 0 | 0 |
| *Gm37800* | 0.431 | 0.001 | 2.659798 | 0 | 0 |
| *Il4i1* | 0.638 | 0.001 | 2.479958 | 0 | 0 |
| *A230065H16Rik* | 0.603 | 0.001 | 2.220012 | 0 | 0 |
| *Ecel1* | 0.448 | 0 | 1.955678 | 0 | 0 |
| *Cd52* | 0.293 | 0.002 | 1.853236 | 0 | 0 |
| *Cr2* | 0.328 | 0 | 1.72423 | 0 | 0 |
| *Mmp23* | 0.552 | 0.009 | 1.72111 | 0 | 0 |
| *Plekhs1* | 0.586 | 0.002 | 1.688413 | 0 | 0 |
| *Rac2* | 0.724 | 0.007 | 1.66724 | 0 | 0 |
| *Wfdc17* | 0.448 | 0.002 | 1.547491 | 0 | 0 |
| *Ncf1* | 0.655 | 0.001 | 1.532593 | 0 | 0 |
| *Sox8* | 0.5 | 0.005 | 1.417345 | 0 | 0 |
| *Tyrobp* | 0.293 | 0.001 | 1.404512 | 0 | 0 |
| *Cd83* | 0.362 | 0.003 | 1.314965 | 0 | 0 |
| *Csf2rb* | 0.31 | 0.001 | 1.312378 | 0 | 0 |
| *AA467197* | 0.397 | 0.002 | 1.262024 | 0 | 0 |
| *Cadm3* | 0.483 | 0 | 1.251538 | 0 | 0 |
| *H2-M2* | 0.345 | 0 | 1.11786 | 0 | 0 |
| *Traf1* | 0.534 | 0.002 | 1.083315 | 0 | 0 |
| *Dsg1a* | 0.362 | 0 | 1.045519 | 0 | 0 |
| *Bcl2a1a* | 0.379 | 0 | 0.986784 | 0 | 0 |
| *Slc2a6* | 0.448 | 0 | 0.963832 | 0 | 0 |
| *Syk* | 0.431 | 0.004 | 0.917025 | 0 | 0 |
| *BC051142* | 0.448 | 0.006 | 0.912784 | 0 | 0 |
| *Bcl2a1d* | 0.517 | 0 | 0.908983 | 0 | 0 |
| *Rassf2* | 0.414 | 0.002 | 0.826474 | 0 | 0 |
| *Csf2rb2* | 0.431 | 0 | 0.796927 | 0 | 0 |
| *Gbp8* | 0.431 | 0.002 | 0.769016 | 0 | 0 |
| *Coro1a* | 0.362 | 0.001 | 0.722442 | 0 | 0 |

| | | | | | |
|---|---|---|---|---|---|
| *Plek* | 0.259 | 0.001 | 0.678191 | 0 | 0 |
| *Laptm5* | 0.276 | 0.002 | 0.671027 | 0 | 0 |
| *Tnfrsf4* | 0.328 | 0.001 | 0.62727 | 0 | 0 |
| *Gbp4* | 0.259 | 0.001 | 0.601246 | 0 | 0 |
| *Cspg4* | 0.259 | 0.001 | 0.588708 | 0 | 0 |
| *Slc16a6* | 0.345 | 0.003 | 0.581011 | 0 | 0 |
| *Gfi1b* | 0.328 | 0.001 | 0.561096 | 0 | 0 |
| *Ecscr* | 0.328 | 0.002 | 0.541479 | 0 | 0 |
| *Napsa* | 0.293 | 0 | 0.50689 | 0 | 0 |
| *Snx20* | 0.259 | 0.002 | 0.493976 | 0 | 0 |
| *Extl1* | 0.259 | 0 | 0.453756 | 0 | 0 |
| *Fxyd5* | 0.259 | 0.003 | 0.733493 | 1.88E-286 | 3.79E-282 |
| *Srgn* | 0.328 | 0.005 | 1.426904 | 3.44E-260 | 6.92E-256 |
| *Ly6i* | 0.69 | 0.023 | 1.917952 | 4.68E-250 | 9.43E-246 |
| *Cfp* | 0.466 | 0.011 | 1.145831 | 1.20E-245 | 2.41E-241 |
| *Syt1* | 0.31 | 0.005 | 0.800201 | 7.14E-221 | 1.44E-216 |
| *Tnfrsf11a* | 0.569 | 0.018 | 1.435914 | 4.71E-215 | 9.47E-211 |
| *Abca13* | 0.293 | 0.005 | 0.638174 | 3.31E-214 | 6.66E-210 |
| *Sh3kbp1* | 0.431 | 0.011 | 0.680126 | 2.27E-204 | 4.57E-200 |
| *Glipr2* | 0.621 | 0.023 | 1.540478 | 8.96E-201 | 1.80E-196 |
| *Gpr137b-ps* | 0.5 | 0.015 | 0.947553 | 1.15E-198 | 2.32E-194 |
| *Tnfaip2* | 0.776 | 0.043 | 2.793595 | 8.97E-173 | 1.80E-168 |
| *Hepacam2.1* | 0.534 | 0.02 | 1.480192 | 1.37E-169 | 2.76E-165 |
| *Slco1a5* | 0.448 | 0.014 | 1.031547 | 3.59E-167 | 7.22E-163 |
| *Vim* | 0.276 | 0.005 | 1.195693 | 9.17E-165 | 1.85E-160 |
| *Cd74* | 1 | 0.081 | 5.11202 | 4.83E-164 | 9.71E-160 |
| *Pde3b* | 0.259 | 0.005 | 0.338614 | 2.01E-157 | 4.04E-153 |
| *Nfkbie* | 0.362 | 0.01 | 0.515248 | 4.51E-153 | 9.08E-149 |
| *Plb1* | 0.397 | 0.013 | 1.194115 | 1.68E-147 | 3.37E-143 |
| *Flt3l* | 0.655 | 0.036 | 1.578435 | 4.52E-143 | 9.10E-139 |
| *Clec12a* | 0.293 | 0.008 | 0.426034 | 1.40E-132 | 2.82E-128 |
| *Rab29* | 0.328 | 0.01 | 0.825804 | 2.06E-125 | 4.14E-121 |
| *H2-Eb1* | 0.483 | 0.023 | 3.628935 | 3.04E-125 | 6.12E-121 |
| *Ncf2* | 0.293 | 0.008 | 0.463194 | 8.12E-123 | 1.63E-118 |
| *Mfhas1* | 0.603 | 0.035 | 1.099555 | 3.90E-122 | 7.86E-118 |
| *Itgb8* | 0.293 | 0.008 | 0.452291 | 3.33E-120 | 6.70E-116 |
| *Rasgef1a* | 0.31 | 0.01 | 0.563651 | 4.01E-115 | 8.06E-111 |
| *Alox12e* | 0.552 | 0.031 | 1.193977 | 7.39E-115 | 1.49E-110 |
| *Gsap* | 0.69 | 0.052 | 1.447816 | 5.57E-110 | 1.12E-105 |

| | | | | | |
|---|---|---|---|---|---|
| *H2-Aa* | 0.517 | 0.03 | 3.987103 | 1.02E-107 | 2.05E-103 |
| *Ero1lb* | 0.534 | 0.031 | 1.22548 | 1.30E-107 | 2.62E-103 |
| *Tnfrsf1b* | 0.431 | 0.021 | 0.648873 | 2.60E-105 | 5.24E-101 |
| *Ccl20* | 0.741 | 0.069 | 6.688476 | 2.54E-102 | 5.12E-98 |
| *Pold1* | 0.431 | 0.022 | 1.201749 | 7.04E-101 | 1.42E-96 |
| *Fabp5* | 0.724 | 0.065 | 3.226804 | 9.78E-101 | 1.97E-96 |
| *Spns2* | 0.328 | 0.013 | 0.60004 | 1.45E-100 | 2.92E-96 |
| *Arhgef9* | 0.276 | 0.009 | 0.523748 | 4.85E-99 | 9.77E-95 |
| *Mia* | 0.397 | 0.019 | 0.535833 | 6.35E-96 | 1.28E-91 |
| *Plekhg1* | 0.345 | 0.016 | 0.75078 | 2.46E-90 | 4.95E-86 |
| *Rab32* | 0.483 | 0.031 | 0.823001 | 1.64E-86 | 3.30E-82 |
| *Kcnj15* | 0.259 | 0.01 | 0.488846 | 3.29E-81 | 6.62E-77 |
| *Birc3* | 0.552 | 0.043 | 0.941839 | 1.18E-80 | 2.37E-76 |
| *H2-Ab1* | 0.534 | 0.045 | 3.684154 | 4.50E-76 | 9.05E-72 |
| *Exoc3l4* | 0.293 | 0.014 | 0.536094 | 2.50E-75 | 5.03E-71 |
| *H2-Q6* | 0.5 | 0.038 | 0.758889 | 1.33E-74 | 2.68E-70 |
| *Pfkfb3* | 0.328 | 0.017 | 0.629849 | 4.77E-74 | 9.59E-70 |
| *Parm1.1* | 0.603 | 0.058 | 1.260557 | 4.80E-73 | 9.66E-69 |
| *Ccdc64* | 0.362 | 0.021 | 0.651153 | 9.99E-73 | 2.01E-68 |
| *9130008F23Rik* | 0.483 | 0.037 | 0.88963 | 4.35E-72 | 8.74E-68 |
| *Cxcr4* | 0.276 | 0.013 | 0.453637 | 5.99E-70 | 1.21E-65 |
| *H2-Q7* | 0.776 | 0.098 | 1.60861 | 1.29E-69 | 2.59E-65 |
| *Gm2a.2* | 0.931 | 0.169 | 2.844312 | 5.82E-69 | 1.17E-64 |
| *Etv3* | 0.776 | 0.105 | 1.543666 | 3.68E-68 | 7.41E-64 |
| *Arhgef40* | 0.259 | 0.012 | 0.459165 | 6.72E-67 | 1.35E-62 |
| *Sirpa* | 0.414 | 0.029 | 0.612285 | 2.29E-66 | 4.61E-62 |
| *Sdhaf1* | 0.569 | 0.054 | 0.753432 | 4.93E-66 | 9.93E-62 |
| *Lpcat4* | 0.397 | 0.029 | 0.812003 | 1.47E-62 | 2.95E-58 |
| *Slc7a11* | 0.759 | 0.108 | 1.763702 | 3.50E-62 | 7.05E-58 |
| *Psmb9* | 0.672 | 0.081 | 0.995676 | 2.12E-61 | 4.27E-57 |
| *Gm15987* | 0.448 | 0.038 | 0.760615 | 4.00E-59 | 8.05E-55 |
| *Slfn2* | 0.379 | 0.028 | 0.799767 | 6.66E-59 | 1.34E-54 |
| *Rasal2* | 0.414 | 0.034 | 0.695616 | 5.82E-58 | 1.17E-53 |
| *Map3k14* | 0.5 | 0.048 | 0.672176 | 1.72E-57 | 3.47E-53 |
| *Nfkb2* | 0.69 | 0.095 | 1.054042 | 1.12E-55 | 2.26E-51 |
| *Tnip1* | 0.638 | 0.08 | 0.982493 | 2.46E-55 | 4.96E-51 |
| *Ctss* | 0.276 | 0.016 | 0.776557 | 6.00E-55 | 1.21E-50 |
| *H2-DMa* | 0.328 | 0.023 | 1.266341 | 8.61E-54 | 1.73E-49 |
| *Lpcat1* | 0.397 | 0.033 | 0.560929 | 1.01E-53 | 2.04E-49 |

| | | | | | |
|---|---|---|---|---|---|
| *Gp2.1* | 0.328 | 0.024 | 2.858139 | 7.06E-53 | 1.42E-48 |
| *Tbccd1* | 0.328 | 0.023 | 0.487215 | 8.24E-53 | 1.66E-48 |
| *Shtn1* | 0.569 | 0.069 | 0.77051 | 2.41E-50 | 4.86E-46 |
| *Samhd1* | 0.672 | 0.104 | 1.358332 | 1.45E-49 | 2.91E-45 |
| *Pde4b* | 0.414 | 0.04 | 0.628461 | 1.15E-47 | 2.32E-43 |
| *Gbp3* | 0.31 | 0.023 | 0.429623 | 2.57E-47 | 5.18E-43 |
| *Dgat2* | 0.345 | 0.028 | 0.541125 | 5.78E-47 | 1.16E-42 |
| *N4bp2l1* | 0.397 | 0.037 | 0.59131 | 7.23E-47 | 1.46E-42 |
| *Mthfsl* | 0.534 | 0.072 | 1.052589 | 7.02E-45 | 1.41E-40 |
| *Psmb8* | 0.914 | 0.211 | 1.292413 | 1.93E-44 | 3.88E-40 |
| *Oasl2* | 0.81 | 0.176 | 1.719944 | 3.02E-44 | 6.08E-40 |
| *H2-DMb1* | 0.379 | 0.037 | 1.007101 | 1.10E-43 | 2.22E-39 |
| *Dclk1* | 0.31 | 0.025 | 0.302787 | 1.18E-43 | 2.38E-39 |
| *Fam20c* | 0.362 | 0.033 | 0.609438 | 3.20E-43 | 6.43E-39 |
| *Nipal1* | 0.5 | 0.064 | 0.89003 | 4.35E-43 | 8.75E-39 |
| *Sema4c* | 0.276 | 0.02 | 0.453447 | 7.59E-43 | 1.53E-38 |
| *AW112010.2* | 0.828 | 0.219 | 5.566619 | 1.08E-42 | 2.18E-38 |
| *Ddx58* | 0.466 | 0.056 | 0.721399 | 3.44E-42 | 6.91E-38 |
| *Calcb* | 0.569 | 0.086 | 2.795794 | 5.94E-42 | 1.19E-37 |
| *Cyba.2* | 0.983 | 0.386 | 3.094591 | 8.65E-42 | 1.74E-37 |
| *Map1b* | 0.517 | 0.069 | 1.063622 | 1.08E-41 | 2.16E-37 |
| *Rhbdl2* | 0.259 | 0.018 | 0.455158 | 1.20E-41 | 2.42E-37 |
| *Vamp5* | 0.759 | 0.166 | 1.591686 | 2.46E-41 | 4.95E-37 |
| *Il4ra* | 0.517 | 0.068 | 0.680346 | 7.95E-41 | 1.60E-36 |
| *Arhgdib* | 0.586 | 0.088 | 0.983278 | 1.32E-40 | 2.66E-36 |
| *Erich3* | 0.31 | 0.026 | 0.398826 | 7.57E-40 | 1.52E-35 |
| *Tnfaip3* | 0.586 | 0.094 | 1.191885 | 1.78E-39 | 3.59E-35 |
| *AI413582* | 0.707 | 0.139 | 1.203036 | 4.58E-39 | 9.21E-35 |
| *Sdcbp2* | 0.655 | 0.121 | 1.508453 | 1.10E-38 | 2.22E-34 |
| *Pard3b* | 0.466 | 0.06 | 0.680274 | 2.14E-38 | 4.31E-34 |
| *Hipk2* | 0.448 | 0.055 | 0.598645 | 2.32E-38 | 4.66E-34 |
| *2310030G06Rik* | 0.586 | 0.098 | 1.033969 | 2.82E-38 | 5.68E-34 |
| *Alox5ap* | 0.31 | 0.029 | 0.810636 | 3.47E-38 | 6.98E-34 |
| *Cldn10.2* | 0.776 | 0.193 | 2.154144 | 3.64E-38 | 7.33E-34 |
| *Sec14l2* | 0.31 | 0.029 | 0.437499 | 5.95E-38 | 1.20E-33 |
| *Dtx3l* | 0.483 | 0.067 | 0.819873 | 1.84E-37 | 3.70E-33 |
| *Sgpp1* | 0.552 | 0.094 | 1.37171 | 2.47E-37 | 4.96E-33 |
| *Pde4dip* | 0.397 | 0.047 | 0.658503 | 9.36E-37 | 1.88E-32 |
| *Mmp15* | 0.534 | 0.084 | 0.825344 | 1.40E-35 | 2.82E-31 |

| | | | | | |
|---|---|---|---|---|---|
| *Relb* | 0.759 | 0.175 | 1.200343 | 2.19E-35 | 4.40E-31 |
| *Ifngr2* | 0.586 | 0.102 | 0.957249 | 2.54E-35 | 5.11E-31 |
| *Atox1* | 1 | 0.815 | 2.472891 | 4.50E-34 | 9.06E-30 |
| *Reep6* | 0.603 | 0.113 | 1.003168 | 5.12E-34 | 1.03E-29 |
| *Fgd2* | 0.259 | 0.022 | 0.295327 | 5.74E-34 | 1.15E-29 |
| *Sppl2a* | 0.707 | 0.158 | 1.186667 | 6.89E-34 | 1.39E-29 |
| *Clmn.1* | 0.483 | 0.071 | 0.711897 | 1.16E-33 | 2.33E-29 |
| *Bcl9* | 0.414 | 0.053 | 0.595308 | 1.46E-33 | 2.93E-29 |
| *Arl5a* | 0.448 | 0.062 | 0.533976 | 1.86E-33 | 3.75E-29 |
| *Lcp1.1* | 0.81 | 0.208 | 1.100745 | 3.90E-33 | 7.85E-29 |
| *Cytip* | 0.259 | 0.024 | 1.324035 | 4.72E-33 | 9.49E-29 |
| *Tank* | 0.569 | 0.099 | 0.749144 | 5.65E-33 | 1.14E-28 |
| *Slc41a1* | 0.362 | 0.043 | 0.507958 | 9.55E-33 | 1.92E-28 |
| *R3hdm1* | 0.638 | 0.117 | 0.715731 | 1.02E-32 | 2.06E-28 |
| *BC021614* | 0.328 | 0.037 | 0.612483 | 3.40E-32 | 6.83E-28 |
| *Dusp4* | 0.483 | 0.076 | 0.934509 | 8.50E-32 | 1.71E-27 |
| *Gda* | 0.431 | 0.06 | 0.617275 | 9.95E-32 | 2.00E-27 |
| *Gpcpd1.1* | 0.707 | 0.161 | 0.973872 | 1.21E-31 | 2.43E-27 |
| *Lrrk1* | 0.397 | 0.052 | 0.489718 | 1.31E-31 | 2.63E-27 |
| *Gpr137b* | 0.707 | 0.167 | 1.108002 | 7.50E-31 | 1.51E-26 |
| *Arsb* | 0.5 | 0.082 | 0.7366 | 1.03E-30 | 2.07E-26 |
| *Tnip3* | 0.362 | 0.046 | 0.858671 | 1.46E-30 | 2.94E-26 |
| *Akt3* | 0.259 | 0.024 | 0.315064 | 1.86E-30 | 3.74E-26 |
| *B2m.1* | 1 | 0.848 | 2.714924 | 2.63E-30 | 5.29E-26 |
| *Pigr.2* | 0.862 | 0.294 | 2.569546 | 3.32E-30 | 6.68E-26 |
| *Rab20* | 0.483 | 0.08 | 0.608772 | 8.27E-29 | 1.66E-24 |
| *Cst3.3* | 1 | 0.84 | 2.697949 | 1.93E-28 | 3.88E-24 |
| *H2afz.3* | 0.983 | 0.849 | 2.266824 | 6.14E-28 | 1.24E-23 |
| *Efhd2* | 0.793 | 0.249 | 1.362423 | 7.23E-28 | 1.45E-23 |
| *Dapk2* | 0.569 | 0.115 | 0.910948 | 8.77E-28 | 1.77E-23 |
| *Klhdc8a* | 0.586 | 0.12 | 0.817749 | 9.37E-28 | 1.89E-23 |
| *Slco4c1* | 0.431 | 0.07 | 0.711548 | 1.14E-27 | 2.30E-23 |
| *Bspry.2* | 0.776 | 0.239 | 1.25903 | 1.30E-27 | 2.62E-23 |
| *Serpinb1a.1* | 0.776 | 0.293 | 3.78124 | 1.91E-27 | 3.84E-23 |
| *Mrpl38* | 0.741 | 0.207 | 1.301419 | 2.03E-27 | 4.09E-23 |
| *Arpc1b.2* | 0.983 | 0.735 | 1.873261 | 2.77E-27 | 5.58E-23 |
| *Ift57* | 0.655 | 0.153 | 0.991368 | 3.37E-27 | 6.77E-23 |
| *Lbh* | 0.397 | 0.06 | 0.717164 | 6.00E-27 | 1.21E-22 |
| *Nfe2l3* | 0.448 | 0.075 | 0.57461 | 1.09E-26 | 2.20E-22 |

| | | | | | |
|---|---|---|---|---|---|
| *Klk11* | 0.276 | 0.031 | 0.376171 | 1.34E-26 | 2.69E-22 |
| *Cxcl16.2* | 0.983 | 0.499 | 1.481739 | 1.39E-26 | 2.80E-22 |
| *Agtrap* | 0.5 | 0.094 | 0.647302 | 2.72E-26 | 5.47E-22 |
| *Tnfrsf11b* | 0.328 | 0.043 | 0.74829 | 3.87E-26 | 7.79E-22 |
| *Nat6* | 0.431 | 0.07 | 0.537633 | 5.26E-26 | 1.06E-21 |
| *Tap2* | 0.672 | 0.169 | 0.992332 | 8.01E-26 | 1.61E-21 |
| *Npc2.2* | 1 | 0.651 | 1.552267 | 8.51E-26 | 1.71E-21 |
| *Msln.3* | 0.948 | 0.503 | 2.66249 | 2.14E-25 | 4.31E-21 |

**Figure E1**

**A**

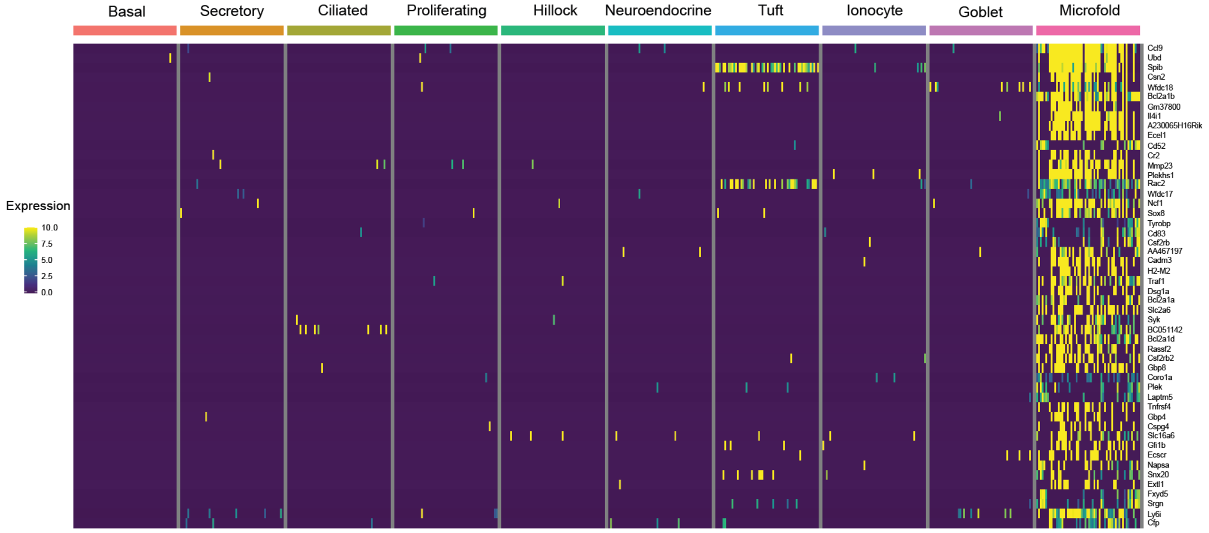
Basal
Secretory
Ciliated
Proliferating
Hillock
Neuroendocrine
Tuft
Ionocyte
Goblet
Microfold
Expression
10.0
7.5
5.0
2.5
0.0
Ccl9
Ubd
Spib
Csn2
Wfdc18
Bcl2a1b
Gm37800
Il4i1
A230065H16Rik
Ecel1
Cd52
Cr2
Mmp23
Plekhs1
Rac2
Wfdc17
Ncf1
Sox8
Tyrobp
Cd83
Csf2rb
AA467197
Cadm3
H2-M2
Traf1
Dsg1a
Bcl2a1a
Slc2a6
Syk
BC051142
Bcl2a1d
Rassf2
Csf2rb2
Gbp8
Coro1a
Plek
Laptm5
Tnfrsf4
Gbp4
Cspg4
Slc16a6
Gfi1b
Ecscr
Napsa
Snx20
Extl1
Fxyd5
Srgn
Ly6i
Cfp

**B**

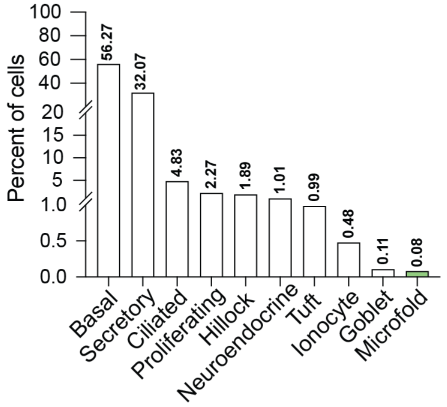
Percent of cells
100
80
60
40
20
15
10
5
1.0
0.5
0.0
56.27
32.07
4.83
2.27
1.89
1.01
0.99
0.48
0.11
0.08
Basal
Secretory
Ciliated
Proliferating
Hillock
Neuroendocrine
Tuft
Ionocyte
Goblet
Microfold
